# Do Motoneurons’ Discharge Rate Slow With Aging? A Systematic Review And Meta-Analysis

**DOI:** 10.1101/2020.12.15.422812

**Authors:** Lucas B. R. Orssatto, David N. Borg, Linda Pendrith, Anthony J. Blazevich, Anthony J. Shield, Gabriel S. Trajano

## Abstract

Nervous system maladaptation is linked to the loss of maximal strength production and motor control with aging. Motor unit discharge rates are a critical determinant of force production; thus, lower discharge rates could be a mechanism underpinning maximal strength and motor control losses during aging. This meta-analysis summarized the findings of studies comparing motor unit discharge rates between young and older adults, and examined the effect of distinct muscles and contraction intensities on the magnitude of discharge rates difference between these two groups. Eligible studies were combined in a meta-analysis, including tested contraction intensities and muscles in different levels, to investigate whether there were differences in discharge rates between younger and older adults. Motor unit discharge rates were higher in younger adults compared to older adults, with a pooled standardized mean difference (SMD) for all studies of 0.63 (95%CI= 0.27 to 0.99). Contraction intensity had a significant effect on the pooled SMD, with a 1% increase in intensity associated with a 0.009 (95%CI= 0.003 to 0.015) change in the pooled SMD. These findings suggest that the reductions in motor unit discharge rates, especially at higher contraction intensities, may be an important mechanism underpinning age-related losses in maximal strength production.

## 1. Introduction

Aging is accompanied by a notable reduction in the ability to produce muscular force and power as well as a decline in the maximal rate of force development, which influences the capacity to perform physical activities of daily living (Bergland and Strand, 2019; Orssatto et al., 2020; Suetta et al., 2019; Tomás et al., 2018). These reductions are clearly observed after 50-60 years of age and the force, power and functional capacity losses accelerate during the subsequent decades of life (Larsson et al., 2019; Suetta et al., 2019; Vandervoort, 2002). Changes within the neuromuscular system have been investigated in order to understand the mechanisms underpinning the differences in force production across the lifespan (Hunter et al., 2016; Larsson et al., 2019; Manini et al., 2013; Orssatto et al., 2018). In addition to alterations within the muscles themselves, the role of neural factors needs to be clearly defined, given that muscle output will be limited by the neural input arriving from the motoneurons.

Both the number of recruited motor units and their discharge rates can be readily modulated by the central nervous system in order to regulate muscle forces. Higher forces are achieved by an increase in the number of active motor units and/or their discharge rates (Enoka and Duchateau, 2017). The relative importance of motor unit recruitment versus discharge rate to the force rise differs significantly between muscles (De Luca and Kline, 2012) and important changes have been observed at the motor unit level during aging (Hassan et al., 2021; Manini et al., 2013; Orssatto et al., 2021a). Over at least the last four decades, researchers have investigated the effects of human aging on motor unit discharge rates during sustained isometric contractions (Kamen et al., 1995; Kirk et al., 2016; Mota et al., 2020; Nelson et al., 1983; Vaillancourt et al., 2003). However, inconsistent results have been observed when comparing young and older individuals, with some studies identifying lower (Christie and Kamen, 2010; Connelly et al., 1999; Kamen and Knight, 2004; Kirk et al., 2019, 2018; Mathew Piasecki et al., 2016a) but others similar mean discharge rates for a given proportional level of force during sustained isometric contractions (Christie and Kamen, 2009; Kallio et al., 2010; Kamen and Roy, 2000; Kirk et al., 2016; Power et al., 2012, 2010). This inconsistency may be partly explained by the wide range of methods used across studies, with variations in the (i) muscles studied; (ii) contraction intensities used (2.5 to 100% of maximal voluntary force); and (iii) small sample sizes (ranging from 5 to 49 young and 5 to 36 older adults). Because of these differences, it has been difficult to draw broad conclusions based on data from any single study. In this case, a systematic review and analysis that pools the available data presented across studies could provide evidence that is more robust in relation to the effect of aging on motor unit discharge rates with special attention to the effects of muscle and contraction intensity.

Given the above, the aim of the present systematic review and meta-analysis with meta-regression was to identify and summarize the findings, and then estimate the effects, of studies comparing motor unit discharge rates between young and older adults during maximal and submaximal isometric sustained contractions. Thereafter, we discuss the potential influence of muscle and contraction intensity on differences in motor unit discharge rates between younger and older individuals.

## 2. Methods

### 2.1. Systematic search strategy

A systematic literature search was conducted in three electronic databases (i.e., PubMed, Web of Sciences, and Scopus) in October 2021. The chosen search terms related to aging (e.g., age, elderly, older, and aging), motor units, and discharge rate (e.g., discharge rate, firing frequency, and inter-spike interval) (Supplementary Material 1). The reference lists of the selected studies were screened for additional studies. The search procedures are reported in the Preferred Reporting Items for Systematic Reviews and Meta-Analyses (PRISMA) flow diagram (Liberati et al., 2009; Moher et al., 2009) (Figure 1). The systematic search strategy was designed by LBRO and GST and conducted independently by LBRO and LP.

**Figure 1.**
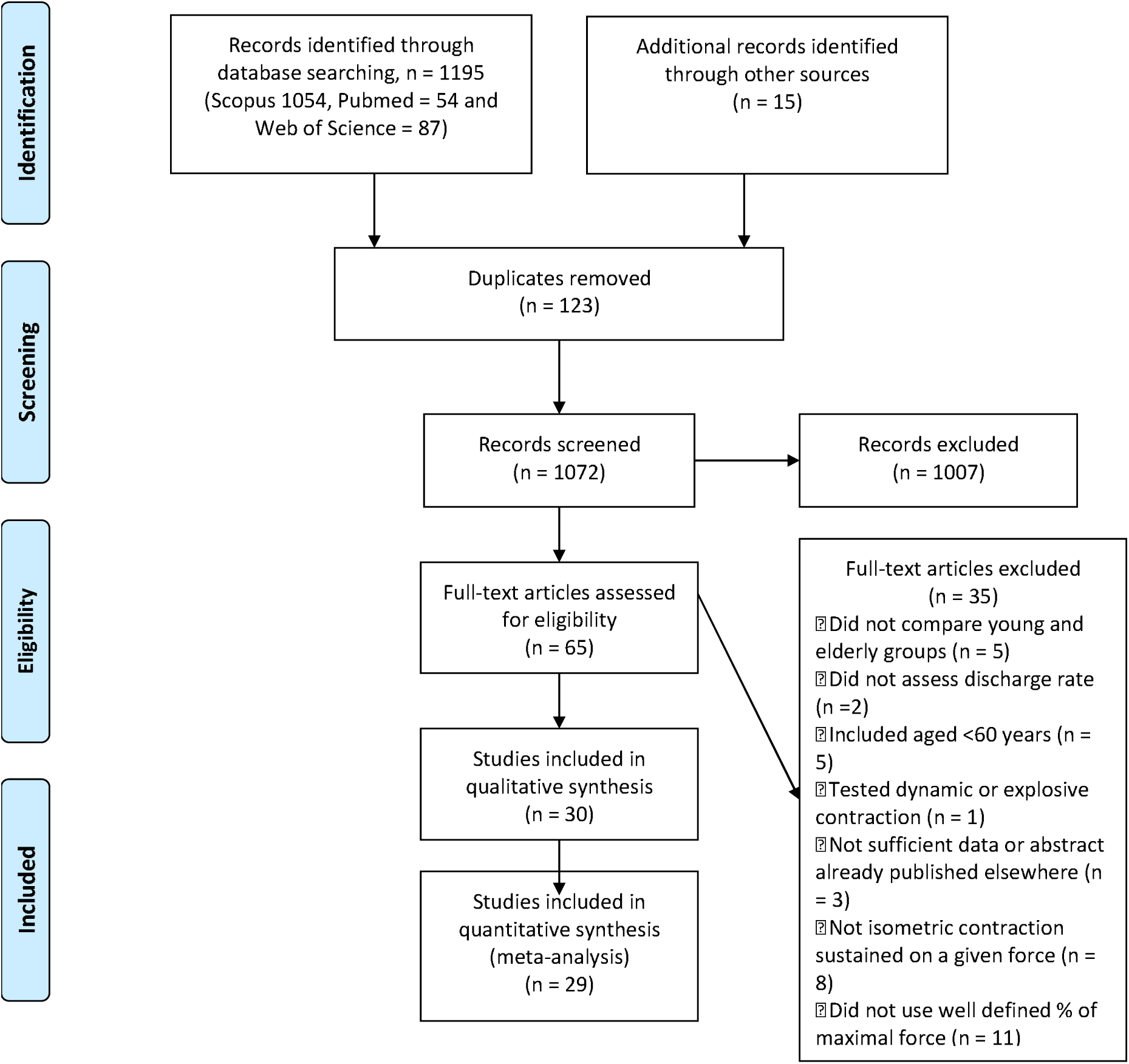
PRISMA flow diagram for the systematic search.

### 2.2. Eligibility criteria

Studies were selected according to the following inclusion criteria: i) compared groups based on age (young = 18–40 years and elderly ≥60 years); ii) adopted electromyographic methods allowing the identification of motor unit discharge rate; iii) discharge rate was measured during a sustained isometric contraction; iv) force level chosen for testing was based on a percentage of each participant’s maximal voluntary force; and v) peer-reviewed and published in English. Studies were excluded if they included participants with neurological (e.g., Parkinson disease, Alzheimer’s) or musculoskeletal disorders (e.g., osteoarthritis, limb injury), involved explosive or dynamic contractions, or if data analyses or the test task were not clearly reported.

### 2.3. Data extraction

Data extracted from the selected studies included the country of the corresponding author, participant sample size, participants’ characteristics and ages, muscles tested, relative force level reached during testing, and mean discharge rates for both young and older groups. For studies reporting chronic or acute intervention results (e.g., exercise training, fatiguing protocol, or visual gain) the baseline or control group/condition data were extracted. Data were converted to discharge rates for studies reporting inter-spike intervals. When data were reported in graphs, WebPlot Digitizer software (version 4.1) was used to extract the data. LBRO and LP conducted all data extraction independently before comparing results; mean variation between researchers was <1%. The mean values from data extracted by LBRO and LP were used for the analyses.

### 2.4. Data analyzes

Our primary interest was to investigate whether there were differences in discharge rates between younger and older adults, with a specific focus on differences in various muscle groups. We were also interested in determining the effect of contraction intensity on differences in motor unit discharge rates between younger and older adults.

Estimates from studies were combined in the meta-analysis using a random effects model (restricted maximum likelihood) and presented as forest plots. We used the standardized mean difference (SMD; Hedges’ *g*) in motor unit discharge rates between younger and older adults, as it was not always clear whether discharge rates were reported per motor unit or per participant. A two-level hierarchical model was fitted because most studies reported several estimates for the same muscle group (i.e., at different contraction intensities). This modelling approach accounts for the clustering of multiple measurements on the same group of subjects (Assink and Wibbelink, 2016). The model assumes that at the first level, individual results in each study are distributed around a study-level effect mean, and that at the second level, the study means are distributed around an overall effect mean. To address our second aim, muscle and contraction intensity were included as moderators in the hierarchical model, to determine their effect on the pooled SMD.

The variance of (the distribution of) the *true* effect sizes (τ^2^) was used as an indicator of between-study heterogeneity, with higher values denoting greater heterogeneity (Rücker et al., 2008). Publication bias was evaluated using a funnel plot (Egger et al., 1997). A sensitivity analysis was performed to determine the robustness of meta-estimates by comparing the pooled SMD and between-study heterogeneity before and after removing the largest and smallest effects, respectively. All analyses were conducted in R (version 4.0.3) using the RStudio environment (version 1.1.447) and the *meta* (Balduzzi et al., 2019), *metafor* (Viechtbauer, 2010) and *dmetar* (Harrer et al., 2019) packages. The α for all tests was set at 5%. The data and R code that support the findings in this article can be accessed at https://github.com/orssatto/SRMA_MU.

## 3. Results

### 3.1. Systematic search

The systematic search conducted in October 2021. It was retrieved 1195 studies and after duplicates removed, 1072 study titles and abstracts were read for eligibility. After removing 1007 studies, 65 studies were read in full and 35 were excluded according to the set criteria, resulting in 30 studies being included in the review. Figure 1 shows the PRISMA flow diagram for all steps of the systematic search.

### 3.2. Study characteristics

Of the 30 included studies, 16 were conducted in the United States of America, 11 in Canada, 1 in Finland, and 2 in the United Kingdom. Studies tested a total of 353 young and 370 older participants. Different muscles were tested, including upper and lower body, flexors and extensors, and hand muscles. Most studies used intramuscular electromyography, except for four studies that used multichannel surface electromyography (5-pin electrodes, n = 2; and 32- or 64-channel electrodes, n = 2). The contraction intensities ranged from 2.5% to 100% of the participants’ maximal voluntary contraction forces.

### 3.3. Meta-estimates

#### 3.3.1. Overall pooled effect

Twenty-nine studies were used for the meta-analysis, in which Kamen and Knight (2004) was not included in the meta-analysis, because we were unable to retrieve all necessary data. The pooled effects for each muscle subgroup of the 29 studies are shown in Figure 2 for lower-body muscles and 3 for upper-body muscles. Motor unit discharge rates were higher in younger adults compared to older adults, with an overall pooled SMD for all studies of 0.63 (95% CI = 0.27 to 0.99, *p* < .001). There was evidence of between-study heterogeneity (τ^2^ = 0.721, 95% CI = 0.388 to 1.393), justifying the use of a random effects model.

**Figure 2.**
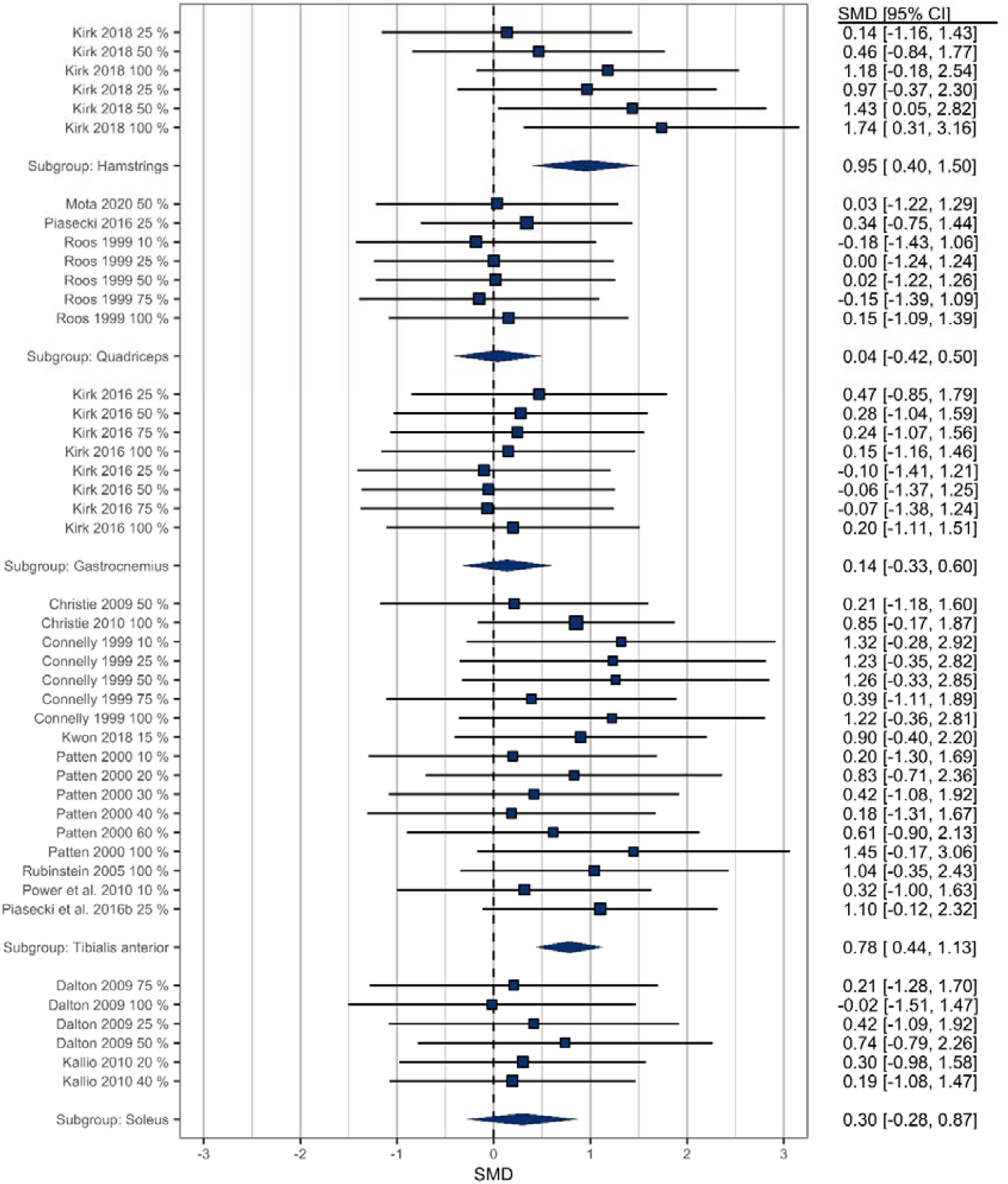
Estimates of differences in motor unit discharge rates between younger and older adults in lower body muscles. Positive estimates denote higher motor unit discharge rates for younger adults and negative estimates denote higher discharge rates for older adults. Note that studies ID are repeated accordingly to the contraction intensity adopted by them. This has been taken into account in the model, preventing the same of the unit-of-analysis problem. SMD, standardized mean difference; 95%CI, 95% confidence interval lower and upper limits.

#### 3.3.2. Effect of muscle and contraction intensity

The test of moderators was significant when both muscle (df = 12) and contraction intensity (df = 78) were included in the model, *F* = 2.482, *p* = .008. There was evidence of an effect of contraction intensity on the pooled SMD, with a 1% increase in contraction intensity associated with a 0.009 (95% CI = 0.003 to 0.015, *p* = .005) change in the SMD. Discharge rates were higher for younger adults in the muscles: *trapezius, biceps brachii*, and *triceps brachii* (Figure 3), and *hamstrings* and *tibialis anterior* (Figure 2). However, these results, along with the absences of differences in other muscles, should be interpreted with caution. Some muscle groups had as few as three estimates available, sometimes came from different intensities from a single study (e.g., *trapezius*). As such, the failure to find any effect is likely the result of very low statistical power. The *first dorsal interosseous* was the only muscle where there may be no difference between younger and older adults, as the estimates came from six different studies. However, the 95% CI is compatible with small negative and small positive effects, and greater precision on this internal is likely required to conclude that discharge rates are not different between younger and older adults in this muscle.

**Figure 3.**
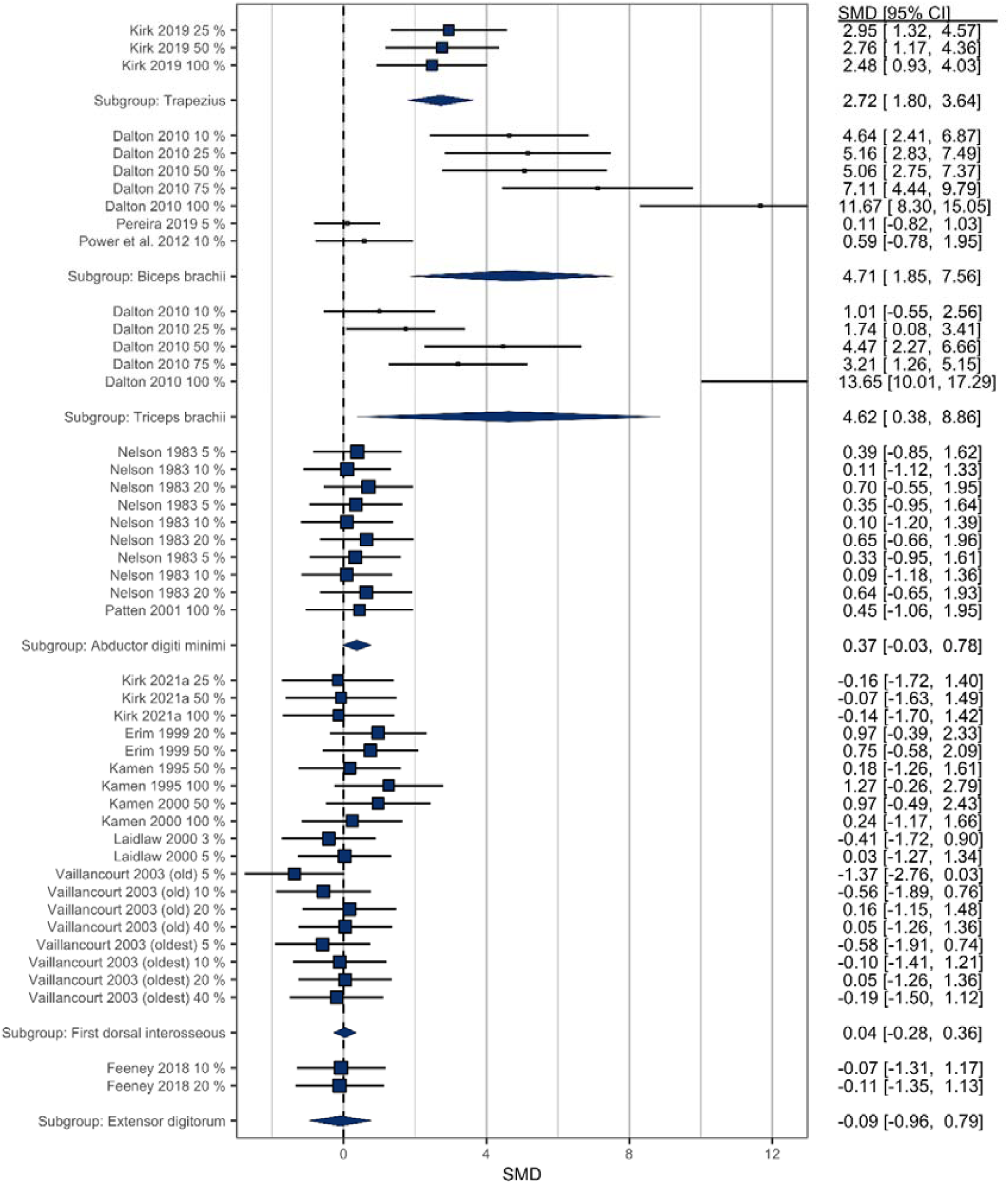
Estimates of differences in motor unit discharge rates between younger and older adults in upper body muscles. Positive estimates denote higher motor unit discharge rates for younger adults and negative estimates denote higher discharge rates for older adults. Note that studies ID are repeated accordingly to the contraction intensity adopted by them. This has been taken into account in the model, preventing the same of the unit-of-analysis problem. SMD, standardized mean difference; 95%CI, 95% confidence interval lower and upper limits.

#### 3.3.3. Publication bias and sensitivity analysis

There was evidence of funnel plot asymmetry (Figure 4A). Removing the study by Dalton et al., (2010) from the meta-analysis removed the asymmetry from the funnel plot (Figure 4B) and led to a marked reduction in the pooled SMD (*g* = 0.47, 95% CI = 0.23 to 0.71, *p* < .001) and a large decrease in between-study heterogeneity (τ^2^ = 0.202, 95% CI = 0.053 to 0.521).

**Figure 4.**
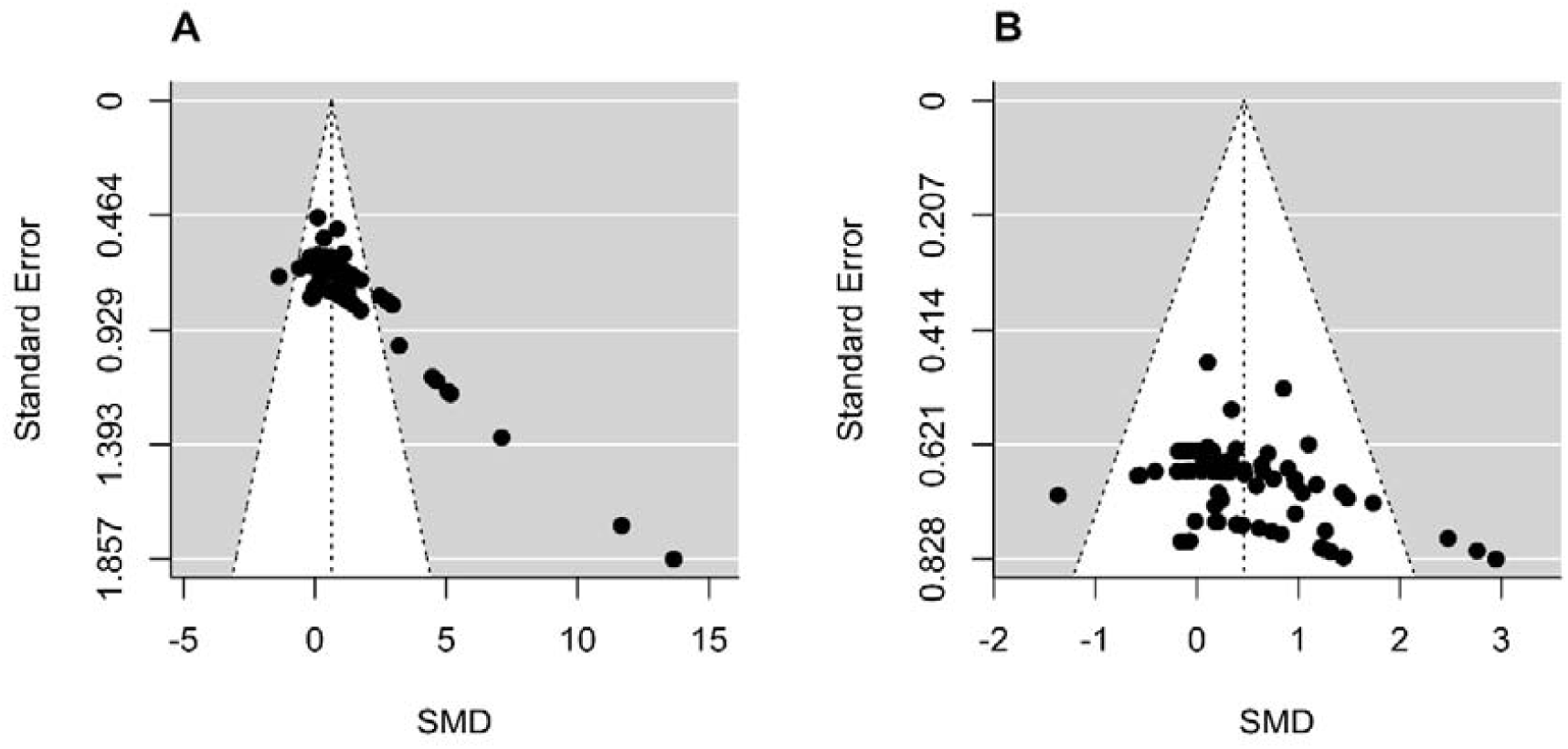
Publication bias funnel plot with all studies (Panel A) and after removing the study by Dalton et al., (2010) (Panel B). Note that Dalton et al., (2010) induced funnel plot asymmetry in Panel A, which has been improved after removing it in Panel B, which led to a marked reduction in the pooled Hedges’g (i.e., SMD).

Removing the smallest effect (*g* = –1.37, *first dorsal interosseous*, 5% contraction intensity (Vaillancourt et al., 2003)) had little influence on both the pooled SMD and between-study heterogeneity (SMD = 0.64, 95% CI = 0.29–0.99; *p* < .001; τ^2^ = 0.701, 95% CI = 0.375–1.359). Removing the largest effect (*g* = 13.65, *triceps brachii*, 100% contraction intensity (Dalton et al., 2010)) had little influence on the pooled SMD, but resulted in a marked reduction in heterogeneity (SMD = 0.62, 95% CI = 0.28–0.95; *p* < .001; τ^2^ = 0.627, 95% CI = 0.328–1.232).

## 4. Discussion

### 4.1. Main findings

The present meta-analytic review interrogated the current literature to determine the effects of aging on motor unit discharge rates. We summarized the findings of studies comparing motor unit discharge rates between young and older adult participants during sustained maximal and submaximal isometric contractions. We also attempted to determine the influence of muscle and contraction intensity on motor unit discharge rates during aging. The main findings were that motor unit discharge rates were higher in young adults (SMD = 0.63), with the 95% confidence interval compatible with small to large effects. Moreover, the magnitude of these differences is greater as contraction intensities increases. It was possible to observe clear differences between young and older adults motor unit discharge rates in *tibialis anterior* and possibly the absence of difference in the *first dorsal interosseous*. The small number of studies testing other specific muscles did not allow to further understand a potential effect of muscle on the differences between young and older adults discharge rates. These results show why inconsistent results have been observed in the extant literature as to the effects of aging on motor unit discharge rates, and they thus contribute to our understanding of the effects of aging on neural control during sustained isometric submaximal and maximal contractions.

### 4.2. Effect of aging and contraction intensity

The reduction in discharge rates, particularly at high contraction levels, might reflect both functional and structural changes within the nervous system during aging. Hypothetically, mechanisms contributing to a potential motor unit discharge rate reduction may include i) a reduced ionotropic input onto the motor unit; ii) the effect of neuromodulation on motoneuron excitability; and iii) the motoneuron structural integrity and motor unit remodeling.

At the motoneuron and motor unit levels, it is also observed several age-related structural and functional changes that may explain the greater reduction in motor unit discharge rate in higher-force contractions. Central to this is that motoneurons are affected by axonal demyelination, atrophy and degeneration during aging (Jang and Remmen, 2011; McKinnon et al., 2015; Misgeld, 2011; Selman et al., 2012). This motoneuron deterioration can trigger muscle fiber denervation, resulting in some fibers being unable to contribute to force production (Aare et al., 2016). Whilst a compensatory mechanism allows low-threshold motoneurons to reinnervate some nearby denervated type II muscle fibers (Hepple and Rice, 2015), it does not prevent the reduction in motor unit discharge rate since the lower-threshold reinnervating motoneuron will exhibit slower discharge rates (Deschenes, 2011; Mathew Piasecki et al., 2016b). As a result, older adults have fewer (McNeil et al., 2005; Mittal and Logmani, 1987; Tomlinson and Irving, 1977) but larger MUs, a greater proportion of which are innervated by lower-threshold motoneurons (Dalton et al., 2008; Lexell and Taylor, 1991). Although debate exists (Piotrkiewicz and Türker, 2017), motor units innervated by higher recruitment-threshold motoneurons should discharge at higher rates during contractions close to maximal force. However, at lower contraction intensities, motor units innervated by lower threshold motoneurons typically discharge at higher rates than those innervated by higher threshold neurons, which typically start to discharge at higher rates as force level increases (Hu et al., 2014; Oya et al., 2009). As older adults typically possess a greater proportion of lower-threshold motor units, this might explain the lower rate relative to younger adults at higher intensities contractions.

The synaptic input onto the motoneurons is another determinant factor to the rate in which a group of motor units discharges. As discussed before, individual motoneurons present a non-linear input-output relation between the received synaptic input and motoneuron discharging due to the presence of persistent inward currents. However, when analyzing a group of motoneurons, it allow the observation of a more linear transformation of common synaptic input into a cumulative train of action potentials to the muscle fibers (Negro and Farina, 2011). Therefore, the discharge output from a group of motoneurons is controlled by receiving ionotropic input in the low-frequency band-width (i.e., effective neural drive) whose gain is adjusted by motoneuronal neuromodulation (Binder et al., 2020; Negro et al., 2009; Orssatto et al., 2021b). The synaptic input onto the motoneurons is decreased with aging (Chase et al., 1985; Maxwell et al., 2018; Rowe et al., 2006), which could be a consequence of reduced intracortical facilitation and increased intracortical inhibition (McGinley et al., 2010; Opie et al., 2020; Todd et al., 2003).

Motor unit discharge rates are also influenced by neuromodulatory inputs (i.e., serotonin and noradrenaline) from the monoaminergic system, which affects motoneuron intrinsic excitability (Binder et al., 2020). Monoamines exert strong facilitation onto motoneurons by triggering persistent inward currents generated by voltage-sensitive sodium and calcium channels. Neuromodulation can change the monoaminergic input onto motoneurons adjusting the amplification of synaptic input according to the motor task demand. When persistent inward current receive a strong monoaminergic drive and is strongly active, the motoneuron intrinsic excitability is increased, allowing it to discharge in higher frequencies (Gorassini et al., 2002; Orssatto et al., 2021b). Therefore, higher intensities contractions presents a greater motoneuronal persistent inward current strength (Orssatto et al., 2021b). In fact, estimates of motoneuronal persistent inward currents have been shown to be reduced in upper and lower limbs of older adults (Hassan et al., 2021; Orssatto et al., 2021a). It has been hypothesized, based in animal and human evidence, that these reductions are mainly explained by the deteriorating age-related adaptations within the monoaminergic system, which indicate reduced noradrenaline and serotonin secretions, and hence input onto motoneurons in older adults (Ko et al., 1997; Liu et al., 2020, 2019; Míguez et al., 1999; Shibata et al., 2006) (See discussions in Orssatto et al., (2021a) and Hassan et al., (2021) for further details). Thus, lower monoaminergic input onto motoneurons would limit its ability to discharge in higher frequencies, especially in higher intensities contractions (Orssatto et al., 2021).

In summary, the lower peak discharge rates observed in older adults could be a result of a reduced synaptic input (descending drive and afferent feedback) received by motoneurons and its capacity to integrate and amplify synaptic input onto discharge rate output via persistent inward currents. This, in addition to the smaller number and proportion of higher-threshold motoneurons may collectively limit motor unit discharge rates during high-intensity, more than low-intensity, contractions.

### 4.3. Muscle subgroups analysis

Age-dependent differences in discharge rates are not readily apparent in the muscle subgroups analyses. Most of these results are underpowered due to the small number of studies and samples sizes for each muscle, except for *tibialis anterior* and *first dorsal interosseous*. Keeping this in mind, the following interpretations should be taken with caution.

The differences observed in *tibialis anterior* (n = 8) and preliminary evidence of differences for *hamstrings* (n = 1) but not in *soleus* (n = 2), *gastrocnemius* (n = 1) and *quadriceps* (n = 3) suggest a pattern of reduction in discharge rates in motor units from lower-limb flexors, rather than from lower-limb extensors. Differences between muscles of different anatomical location might speculatively be explained by the characteristics of the descending tracts that innervate these muscle groups (Taylor et al., 2000). For example, previous studies in the ventral spinal cord of the decerebrate cat and neonatal rat as well as estimations of motoneuron excitability in humans have shown a greater excitability of extensor than flexor motoneurons (Cotel et al., 2009; Hounsgaard et al., 1988; Wilson et al., 2015). Lower body extensors commonly serve an anti-gravity role, meaning that they are active for longer periods and produce greater cumulative force than flexor muscles during activities of daily living, including during upright standing and locomotor propulsion (e.g., walking) (Masani et al., 2013; Soames and Atha, 1981). These functions may be associated with a lesser risk of discharge rate loss with aging. Additional evidence for this assertion is provided by the findings that, while disuse can aggravate the deleterious effects of aging on the nervous system, master-level (i.e., older adult) athletes show a substantial preservation of neural function (Aagaard et al., 2010; Hvid et al., 2018; McGregor et al., 2011; Unhjem et al., 2016). However, further studies testing *hamstrings, soleus, gastrocnemius*, and *quadriceps* should be developed to confirm this hypothesis.

For the upper body muscles, there were no differences between younger and older adults for any muscles responsible for hand movements, such as *first dorsal interosseous* (n = 6), *extensor digitorum* (n = 1), and *abductor digiti minimi* (n = 2). It could be speculated, with caution, that hand muscles are constantly used during the lifespan, which could have prevented discharge rate loss with aging, as discussed before. Conversely, the estimates from *anconeus* (n = 1), *trapezius* (n = 1), *biceps brachii* (n = 3), and *triceps brachii* (n = 1) could suggest lower discharge rates for older adults. Interestingly, the SMD for *triceps brachii* and *biceps brachii* from Dalton et al., (2010) is considerably bigger than the observed difference in other studies and muscles, which noticeably affected the subgroups, overall pooled estimates and unsurprisingly, between-study heterogeneity (See Figure 4). For example, Pereira et al., (2019) and Power et al., (2012) showed a very low and non-significant SMD on *biceps brachii*. Therefore, any specific conclusion regarding these muscles should be avoided.

### 4.4. Strengths and Limitations

Despite the relevance of the current findings, some limitations should be mentioned. First, the small number of studies investigating different specific muscles other than tibialis anterior and *first dorsal interosseous* reduced the statistical power for them. Also, our search only identified studies conducted in more economically developed countries (i.e., USA, Canada, Finland, and United Kingdom). It is unclear if, for example, whether the potentially longer life expectancy and better physical health of individuals from developed countries might influence the research findings. Also, studies investigating sex-related effects of aging on motor unit discharge rates are lacking. Only one study (Pereira et al., 2019) compared older and younger men and women, while other studies only tested men, combined male and female data, or did not report the participants’ sex (Table 1). As such, it was not possible to explore potential sex-related differences on motor unit discharge rate reductions during aging; future studies are clearly warranted to address this issue.

**Table 1.** Studies characteristics

| Authors | Country | Young participants |  | Elderly participants |  | Characteristics | Tested muscles | EMG methods | Contraction intensity (% of MVC) |
| --- | --- | --- | --- | --- | --- | --- | --- | --- | --- |
|  |  | Age (years) | Sample (sex) | Age (years) | Sample (sex) |  |  |  |  |
| Christie and Kamen, 2009 | USA | 23.9 ± 4.1 | 8 (F=4; M=4) | 75.6 ± 6.7 | 8 (F=3; M=5) | Inactive to moderately active (30 min of exercise three times per week or less | Tibialis anterior | Intramuscular | 50% |
| Christie and Kamen, 2010 | USA | 21.9 ± 3.1 | 30 (F=15; M=15) | 72.9 ± 4.6 | 30 (F=15; M=15) | Relatively sedentary (structured physical activity less than 3 days/week). | Tibialis anterior | Intramuscular | 100% |
| Connelly et al., 1999 | Canada | 20.8 ± 0.8 | 6 (M) | 82.0 ± 1.7 | 6 (M) | Recreationally active with a mean of 6.7h/week (old) and 7.6 h/week (young) | Tibialis anterior | Intramuscular | 10, 25 50, 75, and 100% |
| Dalton et al., 2009 | Canada | 23.5 ± 2.9 | 6 (M) | 75.3 ± 4.1 | 6 (M) | Recreationally active. Young: University students. Old: in a program to maintain muscular endurance, cardiovascular fitness, and flexibility. | Soleus | Intramuscular | 25, 50, 75, and 100% |
| Dalton et al., 2010 | Canada | 24 ± 2.4 | 6 (M) | 83 ± 9.8 | 6 (M) | Healthy and recreationally active. | Triceps brachii and biceps brachii | Intramuscular | 10, 25 50, 75, and 100% |
| Erim et al., 1999 | USA | 30.2 ± 5.7 | 10 (NR) | 76.9 ± 6.6 | 10 (NR) | No activity level or hand usage was recorded. | First dorsal interosseous | Intramuscular | 20 and 50% |
| Feeney et al., 2018 | USA | 25 ± 4 | 13<br>(F=8;<br>M=5) | 78 ± 5 | 12<br>(F=5;<br>M=7) | No history of neurological disease or injury to the upper extremity | Extensor digitorum | Surface 32-channels electrode | 10 and 20% |
| Kallio et al., 2010 | Finland | 27.3 ± 3 | 10 (M) | 70.4 ± 5 | 13 (M) | Physically active males without any history of neuromuscular or vascular disease. | Soleus | Intramuscular | 20 and 40% |
| Kamen et al., 1995 | USA | R(21 – 33) | 7 (NR) | R(67 – 85) | 7 (NR) | Free of musculoskeletal and neuromuscular complaints. Varied in fitness levels, to give a representative sample of elderly adults. | First dorsal interosseous | Intramuscular | 50 and 100% |
| Kamen and Roy, 2000 | USA | 27.7 ± 4.3 | 7 (NR) | 74.9 ± 7.1 | 8 (NR) | Free of any known neuromuscular, musculoskeletal or cardiopulmonary disorders. | First dorsal interosseous | Intramuscular | 50 and 100% |
| Kamen and Knight, 2004 | USA | 21<br>(R 18 - 29) | 8 (NR) | 77<br>(R 67 – 81) | 7 (NR) | None of these persons had resistance training experience within the previous year. | Vastus lateralis | Intramuscular | 10 and 50% |
| Kirk et al., 2016 | Canada | 27 ± 3 | 10 (M) | 81 ± 4 | 10 (M) | Free from any known neuromuscular, respiratory, cardiovascular and metabolic illness and none were systematically trained or sedentary. | Gastrocnemius | Intramuscular | 25, 50, 75, and 100% |
| Kirk et al., 2018 | Canada | 26 ± 4 | 11 (M) | 80 ± 5 | 10 (M) | Involved in recreational exercise programs three times a week. | Biceps femoris, semitendinosus and semimembranosus | Intramuscular | 25, 50, and 100% |
| Kirk et al., 2019 | Canada | R(22 – 33) | 10 (M) | R(77 – 88) | 10 (M) | Young participants were normally active University students. The older participants were independently living and participated three times per week in an organized fitness and light exercise program. | Superior trapezius | Intramuscular | 25, 50, and 100% |
| Kirk et al., (2021a) | Canada | 28 ± 5 | 5 (M) | 85 ± 6 | 5 (M) | Free from known neurological, metabolic, or orthopedic disease with no prior injury to the hand or upper limb. | First dorsal interosseous | Intramuscular | 25, 50, and 100% |
| Kirk et al., (2021b) | Canada | 28 ± 5 | 8 (M) | 83 ± 5 | 13 (M) | Free of prior injury to the upper limb and any neurological, metabolic, or orthopedic illness. | Anconeus | Intramuscular | 100% |
| Kwon and Christou, 2018 | USA | 22.6 ± 4.1 | 11<br>(F=6;<br>M=5) | 74.1 ± 5.7 | 12<br>(F=5;<br>M=7) | Healthy without any neurological impairment and were moderately active. | Tibialis anterior | Surface 5-pins electrode | 15% |
| Laidlaw et al., 2000 | USA | 26.6 ± 1.6 | 8<br>(F=4;<br>M=4) | 73.4 ± 6.1 | 14<br>(F=8;<br>M=6) | No known neuromuscular disorders and their vision corrected to normal levels. | First dorsal interosseous | Intramuscular | 3 and 5% |
| Mota et al., 2020 | USA | 25 ± 3 | 12 (M) | 75 ± 8 | 12 (M) | During the 6 months prior to study participation, enrolled participants refrained from lower body resistance training (< three times monthly) or other structured exercise (e.g., jogging) more than 30 min per day, three times per week | Vastus lateralis | Surface 5-pins electrode | 50% |
| Nelson et al., 1983 | USA | 29.6 ± 4.2 | 13 (NR) | 65.5 ± 2.2 | 13 (NR) | No current or history of peripheral nerve dysfunction, no current usage of medication known to affect neuronal conduction, and no indication of muscle weakness | Abductor digiti minimi | Intramuscular | 5, 10, and 20% |
|  |  |  |  | 74.8 ± 2.9 | 9 (NR) |  |  |  |  |
|  |  |  |  | 83.5 ± 2.8 | 10 (NR) |  |  |  |  |
| Patten and Kamen, 2000 | USA | 19.2 ± 1.6 | 6 (F=3; M=3) | 70.3 ± 3.9 | 6 (F=2; M=4) | Regular participation in vigorous physical activity since at least the age of 40 years. Young and older adult subjects were matched for current participation in physical activity | Tibialis Anterior | Intramuscular | 10, 20, 30, 40, 60, and 100% |
| Patten et al., 2001 | USA | 23.2 ± 3.5 | 6 (F=3; M=3) | 75.8 ± 7.4 | 6 (F=3; M=3) | Mean weekly time spent participating in physical activity was similar between young (128 min) and older (138 min) adult groups. | Abductor digiti minimi | Intramuscular | 100% |
| Pereira et al., 2019 | USA | 21.9 ± 2.8* | 49 (F=25; M=24) | 67.7 ± 5.8* | 36 (F=19; M=17) | No statistical differences for physical activity level between young and elderly participants. | Biceps Brachii | Surface 64-channels electrode | 5% |
| Piasecki et al., 2016 | United Kingdom | 25.3 ± 4.8 | 22 (M) | 71.4 (6.2) | 20 (M) | Habitually physically active (not sedentary or participating in competitive sport at regional level or above). | Vastus lateralis | Intramuscular | 25% |
| Piasecki et al., 2016 | United Kingdom | 26 ± 4 | 18 | 71 ± 4 | 14 | From the university population and the local community. All participants were healthy and recreationally active. | Tibialis Anterior | Intramuscular | 25% |
| Power et al., 2010 | Canada | 27 ± 3 | 10 | 66 ± 3 | 10 | Recreationally active young university students, and older adults from a local exercise program designed to maintain cardiorespiratory endurance and flexibility. | Tibialis Anterior | Intramuscular | 10% |
| Power et al., 2012 | Canada | 27 ± 5 | 9 | 70 ± 5 | 9 | Recreationally active young university students and older adults. Older adults participating in non-systematic light walking and calisthenic activities. | Biceps Brachii | Intramuscular | 10% |
| Roos et al., 1999 | Canada | 26.2 ± 4.1 | 13 (M) | 80 ± 5.3 | 12 (M) | Moderately active | Vastus medialis | Intramuscular | 10, 25, 50, 75, and 100% |
| Rubinstein and Kamen, 2005 | USA | 19.2<br>(R 18-30) | 34 (W) | 73.1<br>(R >69) | 12 (W) | Engaged in moderate levels of daily physical activity (3–5 days/week). | Tibialis Anterior | Intramuscular | 100% |
| Vaillancourt et al., 2003 | USA | 22 ± 1 | 10<br>(F=5;<br>M=5) | 67 ± 2 | 10<br>(F=5;<br>M=5) | The subjects were naive to the purpose of the experiment and none had a history of a neurological disorder | First dorsal interosseous | Intramuscular | 5, 10, 20, 40% |
|  |  |  |  | 82 ± 5 | 10<br>(F=5;<br>M=5) |  |  |  |  |

R = range; EMG, electromyography; NR, not reported; M, male; F, female; \*pooled mean ± standard deviation from male and female groups.

Another important limitation of the current literature is mostly adopting sustained isometric contractions in which motor unit discharge rates were measured during the contraction plateau were included in our systematic search. This methodological restriction allowed a better control for between-study comparisons because motor unit discharge behavior is contraction mode dependent (i.e., isometric vs. concentric vs. eccentric) and varies with movement velocity (i.e., fast, or slow) and the phase of isometric force production (i.e., torque rise, plateau, or decline). However, this inclusion criterion could mask some potential age-related differences that may occur in other contractions or movement tasks. For example, evidence exists that motor unit discharge rates during explosive contractions may be more sensitive to aging (Klass et al., 2008). Also, lower motor unit discharge rates were observed for older adults during non-sustained isometric contractions (Hassan et al., 2021; Lucas B.R. Orssatto et al., 2021a). Conversely, dynamic contractions have been shown to present no age-related discharge rates differences compared to isometric contractions (Kirk et al., 2021b). However, more studies are required before clear conclusions can be drawn. Therefore, our findings may only be reflective of those during isometric contractions.

## 5. Conclusions

The present systematic review identified 30 studies reporting contrasting results regarding the differences in motor unit discharge rates, during sustained isometric contractions between young and older adults. In overall, motor unit discharge rates were lower in older adults. Also, the magnitude of differences between young and older adults is clearly augmented as contraction intensity increases. These findings suggest the reductions in motor unit discharge rates as an important mechanism underpinning the age-related losses in maximal strength production and motor function.

The muscle subgroups analysis indicated that discharge rates were lower in older adults for *tibialis anterior, hamstrings, biceps brachii, triceps brachii, and trapezius*. However, *tibialis anterior* was the only muscle where a clear difference was observed and estimates came from several studies, rather than only a single study. The absence of differences for *soleus, gastrocnemius, quadriceps, extensor digitorum*, and *abductor digiti minimi* should also be interpreted with caution since the estimates were obtained from one or two studies only. The *first dorsal interosseous* was the only muscle where no difference was observed, and estimates came from several studies. However, greater precision on the SMD is needed to definitively conclude there is no difference between younger and older adults in this muscle. Nonetheless, this evidences that the differences in motor unit discharge rates should be continuously explored in different muscles along with the mechanisms underpinning these reductions.

It is important to consider that our findings may only be reflective of motor unit behavior during isometric contractions. There was also considerable methodological heterogeneity among studies regarding the muscles examined, contraction intensities used, discharge rate measurement techniques, and the age-range of participants. These aspects of study design require careful consideration by future investigations.

## Notes

https://github.com/orssatto/SRMA_MU

### Competing Interest Statement

The authors have declared no competing interest.

